# Single cell mRNA signals reveal a distinct developmental state of *KMT2A*-rearranged infant B-cell acute lymphoblastic leukemia

**DOI:** 10.1101/2021.12.17.473141

**Authors:** Eleonora Khabirova, Laura Jardine, Tim H. H. Coorens, Simone Webb, Taryn D. Treger, Justin Englebert, Tarryn Porter, Elena Prigmore, Grace Collord, Alice Piapi, Sarah Teichmann, Sarah Inglott, Owen Williams, Olaf Heidenreich, Matthew D. Young, Karin Straathof, Simon Bomken, Jack Bartram, Muzlifah Haniffa, Sam Behjati

## Abstract

Infant B-cell acute lymphoblastic leukemia (B-ALL) has not followed the increasing trend towards cure seen in other childhood B-ALLs. The prognosis for infants with *KMT2A* gene fusions is especially poor, and the origins of this aggressive leukemia remain unknown. Here, we investigated the developmental state of *KMT2A*-rearranged infant B-ALL within hematopoietic hierarchies of human fetal bone marrow, using bulk mRNA meta-analysis of childhood leukemia and examination of single lymphoblast transcriptomes. *KMT2A*-rearranged infant B-ALL was uniquely dominated by an early lymphocyte precursor (ELP) state. Direct comparison of infant lymphoblasts with ELP cells distilled the core oncogenic transcriptome of cancer cells which harboured potentially targetable hybrid myeloid-lymphoid features. Overall our quantitative molecular analyses demonstrate a distinct developmental state of *KMT2A*-rearranged infant B-ALL.

## Main

Once a universally fatal disease, acute B lymphoblastic leukemia (B-ALL) of childhood is curable in the majority of cases. An exception is B-ALL arising in children younger than one year of age (infant B-ALL), which remains fatal in more than 50% of children ^1,2^. Most cases (70-80%) of infant B-ALL are associated with rearrangements of the *KMT2A* gene (encoding a histone methyltransferase), which confers an especially poor prognosis ^2^. Various hypotheses have been proposed to account for the aggressive nature of infant B-ALL. In particular, it has been suggested that infant lymphoblasts retain myeloid features that confer resistance to treatment strategies aimed at ALL ^3^. Disappointingly, whilst protocols incorporating strategies from acute myeloid leukemia (AML) therapy marginally increased survival, additional intensification has not improved this further ^1^. Similarly, salvage treatments which have proven successful in high-risk lymphoblastic leukemias, such as allogeneic stem cell transplantation or chimeric antigen receptor-T cells targeting B-cell antigens, produce disappointing outcomes in infant B-ALL ^4,5^. It is noteworthy that infant B-ALL not associated with *KMT2A* fusion, especially those with *NUTM1* gene rearrangements, confer a more favourable prognosis ^6,7^, and that *KMT2A* rearrangements in the setting of adult B-ALL are also considered high-risk ^8^. These observations raise the question whether the aggressive clinical behaviour of *KMT2A*-rearranged infant B-ALL is underpinned by a distinct cellular phenotype.

Leukemias are primarily classified by their morphological appearance, immunophenotype, as assessed by flow cytometric analyses of key hematopoietic markers, and cytogenetic changes. Generally, this approach is likely to capture the differentiation state of most leukemias accurately. Occasionally, it may be erroneous when cancer cells utilise key hematopoietic genes aberrantly, particularly in leukemias that are driven by mutations in genes that facilitate lineage plasticity, such as *KMT2A*. In this context, a quantitative molecular assessment of hematopoietic cell states that does not rely on any individual marker, but instead builds on entire cellular transcriptomes, would provide an unbiased readout of cell states. Such high-resolution assessments are now feasible using single-cell mRNA sequencing to directly compare cancer cells to normal cells, including to fetal and adult hematopoietic cells ^9–12^. We set out to study the developmental phenotype of *KMT2A*-rearranged infant B-ALL, by comparing cancer to normal human hematopoietic cells.

## Results

### Cell signal analysis of 1,665 leukemia transcriptomes

The starting point of our investigation was a meta-analysis of 1,665 bulk transcriptomes representing the entire spectrum of childhood ALL and AML across two cohorts; St Jude Children’s Research Hospital (n=589) and TARGET (Therapeutically Applicable Research to Generate Effective Treatments; n=1076) **(Fig. 1A; Table S1)**. We determined the predominant hematopoietic cell signal within each bulk leukemia transcriptome by deconvolution. We chose a deconvolution method that uses entire transcriptomes to determine cell signals within bulk mRNA data, and quantify what proportion of the cancer bulk cannot be accounted for by normal reference cells ^13^. As childhood leukemias, and infant ALL in particular, are generally thought to arise *in utero* ^*14,15*^, we applied fetal hematopoietic cells as the reference in our analyses. To this end, we utilised recent single-cell mRNA data from ∼60,000 fetal bone marrow cells, which captured the greatest breadth of fetal hematopoietic cell types to date ^9^ **(Table S2)**. We adopted the annotation of normal cell types directly from the fetal bone marrow data analysis ^9^ and supplemented the hematopoietic reference with a control fetal cell type that should not be present in human leukemia samples – Schwann cell precursors (SCP) derived from human fetal adrenal glands ^16^.

**Fig. 1.**
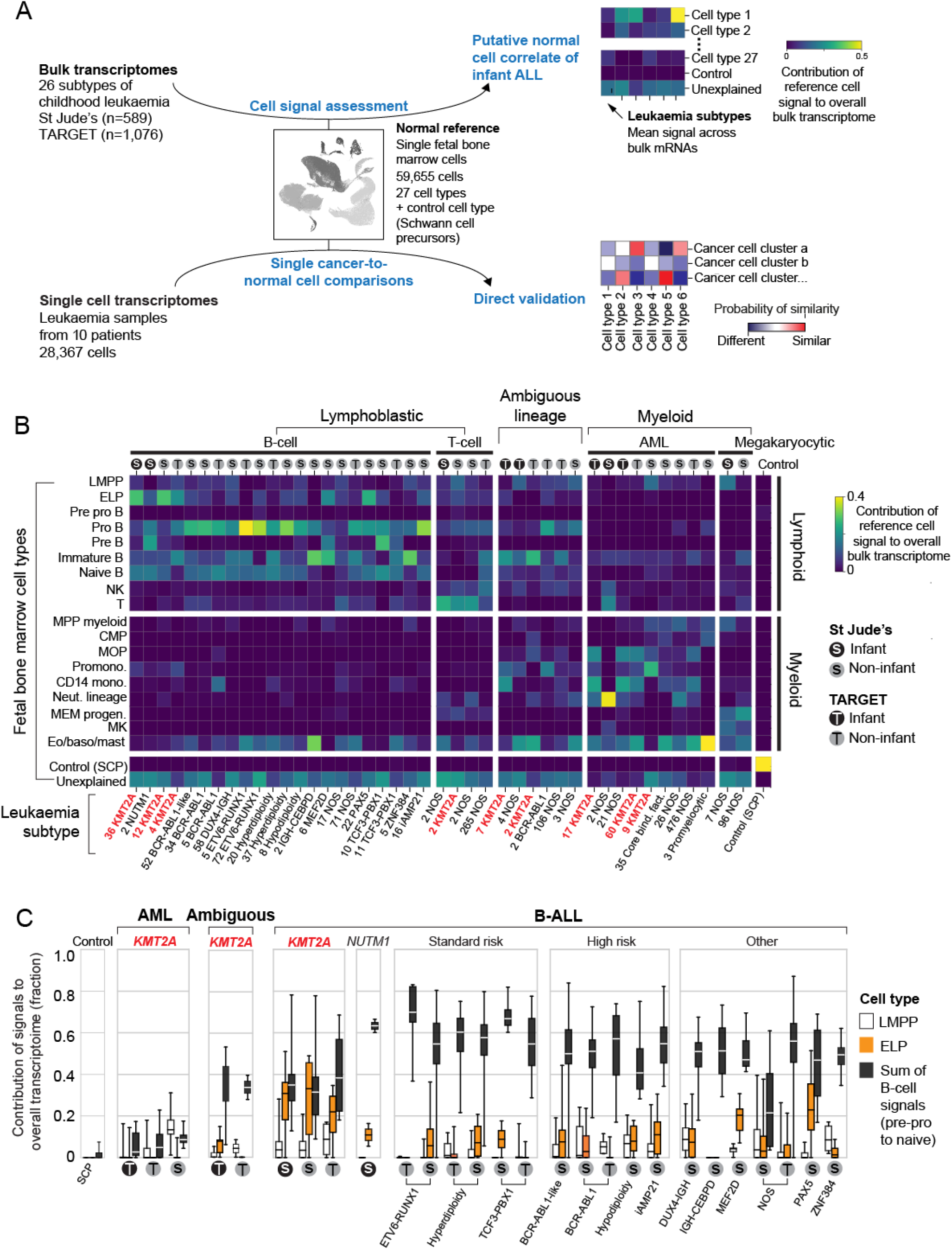
Cell signal analysis of 1,665 leukemia transcriptomes reveals an ELP state in KMT2A-rearranged B-ALL. (**A**) Schematic overview of study approach. We assessed the differentiation state of *KMT2A*-rearranged infant ALL by measuring signals of human fetal bone marrow cell types across the entire spectrum of childhood leukemia in data derived from two different cohorts (St Jude’s, TARGET). We then validated cell signals by single cell mRNA sequencing for direct comparison of cancer and normal cells. (B) Heatmap showing mean cell signals of human fetal bone marrow cells (y-axis) in human leukemia bulk transcriptomes subdivided by genetic subtype (see labels underneath), age (grey circle = infant; black circle = non-infant), and source (S = St Jude’s; T = TARGET). Numbers next to labels refer to case load per subtype. Subtypes with only one case were excluded from analysis. (C) Box and whisker plots showing contributions of signals – LMPP, ELP and latter B-cell stages (i.e. pre-pro B, pro-B, pre-B and naive B combined) to the transcriptome of leukemias (see x-axis labels). Centre line=median, whiskers=min/max and box limits=25%/75% quartiles. Risk refers to the clinical cytogenetic risk as defined in the protocol of the current European ALL trial, “ALLTogether” (EudraCT number: 2018-001795-38).

A global overview of cell signals in bulk childhood leukemia transcriptomes showed expected patterns, namely myeloid signals in myeloid leukemias, T-cell signals in T-ALL and imprints of the various stages of B-cell development in B-ALL **(Fig. 1B)**. Transcriptional signatures from the control SCP population did not contribute to leukemias (negative control analysis) and matched itself (i.e. SCPs) perfectly with no unexplained signal (positive control analysis). *KMT2A*-rearranged infant B-ALL exhibited distinct cell signals with a marked contribution of early lymphocyte precursors (ELP). ELP are oligopotent lymphoid precursors capable of differentiating along different lymphocyte lineages and which retain minimal myeloid differentiation capacity *in vitro* ^*17,18*^. Defined as CD34^+^CD127^+^CD10^−^CD19^-^ cells, they sit upstream of pre-pro B and pro B progenitors in the B lymphopoiesis hierarchy ^18^.

### An early lymphocyte precursor signal in KMT2A-rearranged B-ALL

To further examine the ELP signal in *KMT2A*-driven infant B-ALL, we examined the ratio of the ELP signal over later stages of B-cell development in each leukemia subtype **(Fig. 1C)**. This quantification demonstrated a significant shift towards the ELP state in *KMT2A*-rearranged infant ALL, compared to other high (cytogenetic) risk B-ALL subtypes (p<10^−19^, Student’s two-tailed t-test), standard (cytogenetic) risk subtypes (p< 10^−31^) and currently unstratified subtypes of B-ALL (p<10^−13^) **(Fig. 1C)**. Amongst *KMT2A*-rearranged infant B-ALL, the ELP signal was present irrespective of fusion partners of *KMT2A*, but strongest in cases harbouring the most common *KMT2A* rearrangement ^19^, the *KMT2A-AFF1* (*MLL-AF4*) gene fusion (p<0.01 compared against other fusion partners; Mann-Whitney rank test) (**Fig. S1**). The leukemia with the second highest relative ELP signal was *PAX5*-mutated B-ALL, although the ELP signal there was accompanied by stronger signals from later B cell stages. In contrast to *KMT2A*-rearranged B-ALL, the difference between ELP signals and later B-cell signals was significant in *PAX5*-mutated B-ALL (p<0.01; Wilcoxon signed-rank test). The similarity of cell signals in *PAX5* and *KMT2A* mutant B-ALL may represent the intimate relationship of *KMT2A* and *PAX5* in regulating B-lymphopoiesis ^20^.

Studying the pattern of ELP signal across disease groups, indicated that the signal was specific to *KMT2A*-rearrangements within a B-cell context, but independent of age for three reasons. Firstly, the ELP signal was not universally associated with *KMT2A*-rearrangements; neither myeloid, nor ambiguous lineage leukemias with *KMT2A*-rearrangements harboured appreciable ELP signals. Secondly, the ELP signal was not driven by young age alone, as other infant leukemias (B-ALL, ambiguous lineage leukemia, AML) exhibited no, or only minimal, ELP signals **(Fig. 1C)**. In particular, infant B-ALL with *NUTM1*-rearrangement (which carries a favourable prognosis) exhibited cell signals more reminiscent of standard-risk childhood B-ALL, with a shift away from ELPs towards later B-cell stages. Thirdly, *KMT2A*-rearranged B-ALL of older children did exhibit marked ELP signals akin to infant *KMT2A*-driven B-ALL. Overall, these findings led us to hypothesise that, relative to other B-ALL, *KMT2A*-rearranged B-ALL exhibits a distinct hematopoietic phenotype primarily resembling ELP cells with limited signals of B-cell development.

### Direct single cancer to normal cell comparison

To validate and further explore this proposition, we performed single-cell RNAseq analysis (10x Genomics) on diagnostic specimens from 6 infants with *KMT2A*-rearranged infant B-ALL and compared these to 4 other infant leukemias: *NUTM1*-rearranged B-ALL (n=1); *KMT2A*-rearranged AML (n=1); megakaryoblastic AML (n=1) and *ETV6-RUNX1* B-ALL (a common type of standard risk childhood B-ALL; n=1) (**Table S3**). From these 10 diagnostic leukemia samples, we obtained a total of 28,367 cells, including 22,379 cancer cells that we identified based on gene expression matching patient-specific diagnostic flow cytometric profiles (**Table S4, Fig. S2**). Using a published cell matching method based on logistic regression ^12,16^, we directly compared leukemia transcriptomes with mRNA profiles of human fetal bone marrow cells to determine which normal cell type the cancer cells most closely matched. We found that *KMT2A*-rearranged infant B-lymphoblasts overwhelmingly resembled ELP cells (**Fig. 2A to 2C**). By contrast, non-ELP cell signals predominated in other types of leukemia, precisely as predicted from the initial deconvolution analysis (**Fig. 1B**). In particular, in the aforementioned subtype of infant B-ALL with a favourable prognosis, *NUTM1*-rearranged infant B-ALL, single cell analysis confirmed the shift towards pre-B cell states and away from ELPs.

**Fig. 2.**
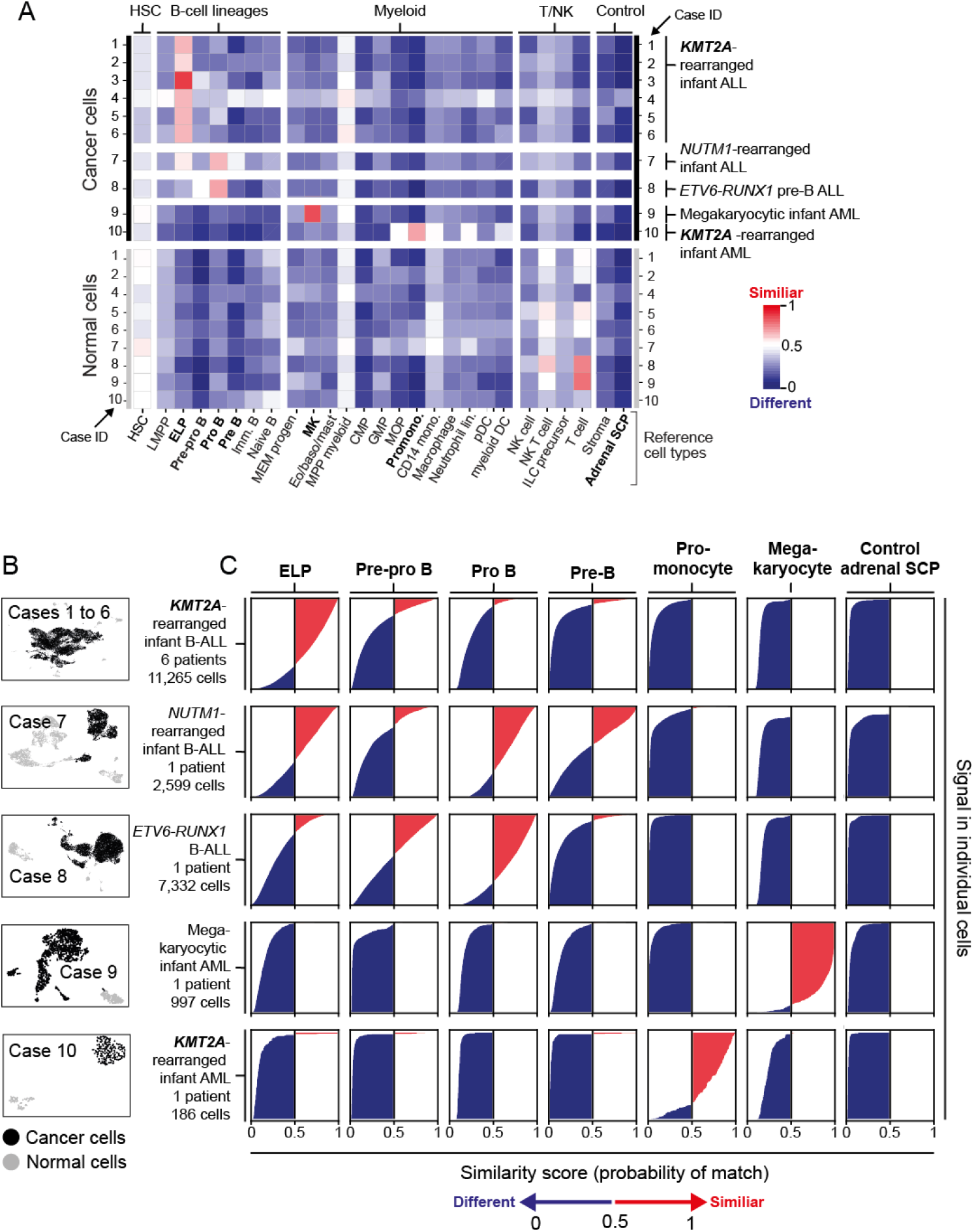
Validation of ELP signals by direct single cancer to normal cell comparison. (**A**) Heatmap showing comparing cell clusters from diagnostic specimens (y-axis) to normal human fetal bone marrow cell types (x-axis; bold labels highlight cell types shown in C). Cell clusters represent cancer (as defined by clinical diagnostic flow cytometric profiles, see Fig. S1) and normal cells of individual patient samples (as per case ID number; see Table S3 for an overview of patients). Heat colors represent the mean probability (across the cell cluster) of a match as determined by logistic regression (red = similar; blue = different). (B) Uniform manifold approximation and projection (UMAP) of single leukemia cell data divided by patient groups and within each heat map by cancer (black dots) and normal cells (grey dots). (C) Logistic regression score of each individual cell against different reference cells per cancer type.

### Phylogenetic timing of the origin of infant ALL

A key question raised by our findings is whether ELPs are the cells of origin of *KMT2A*-rearranged infant B-ALL, or whether leukemia cells arise from another precursor and differentiate/de-differentiate into an ELP-like state at which they arrest. A rare case of lineage switching from *KMT2A*-rearranged infant B-ALL to *KMT2A*-rearranged AML provided the opportunity to directly determine the cell of origin in phylogenetic temporal terms (**Fig. 3A**). We first assessed cell signals in bulk transcriptomes (in replicates) derived from the child’s *KMT2A*-rearranged B-ALL and AML. Once again, we observed that ALL, but not AML transcriptomes, exhibited an ELP signal (**Fig. 3B**). To determine the phylogeny of the cancers, we performed whole genome DNA sequencing of AML, ALL and remission bone marrow, and called all classes of variants, using an extensively validated mutation-calling pipeline ^21^ (variant list in **Table S5**). We determined the phylogeny of each leukemia and of remission bone marrow. The remission sample and leukemias shared two mosaic (early embryonic) base substitutions, representing the first cell divisions of the zygote ^22,23^. Thereafter, normal blood and leukemia lineages diverged. The common leukemia lineage (i.e. mutations shared between ALL and AML, but not the remission sample) comprised only six base substitutions along with the *KMT2A*-rearrangement (**Fig. 3C to 3D**), defining an early developmental window during which the translocation formed. Assuming a mutation rate of at least 0.9 substitutions per cell division, as recently established in human fetal hematopoietic cells ^24^, six substitutions would place the emergence of the *KMT2A*-rearrangement in early embryonic development, before hematopoietic cell specification. After acquisition of the *KMT2A* fusion, the leukemia lineages diverged and gave rise to independent cancers, each exhibiting distinct phenotypes and somatic changes (including point mutations, copy number profiles, mutational signatures) (**Fig. 3C to 3E**). Although this single case may not be representative of infant ALL generally or of lineage switch leukemias specifically, it demonstrates that the transcriptional state of cancer cells cannot unambiguously be used to infer its cell of origin.

**Fig. 3.**
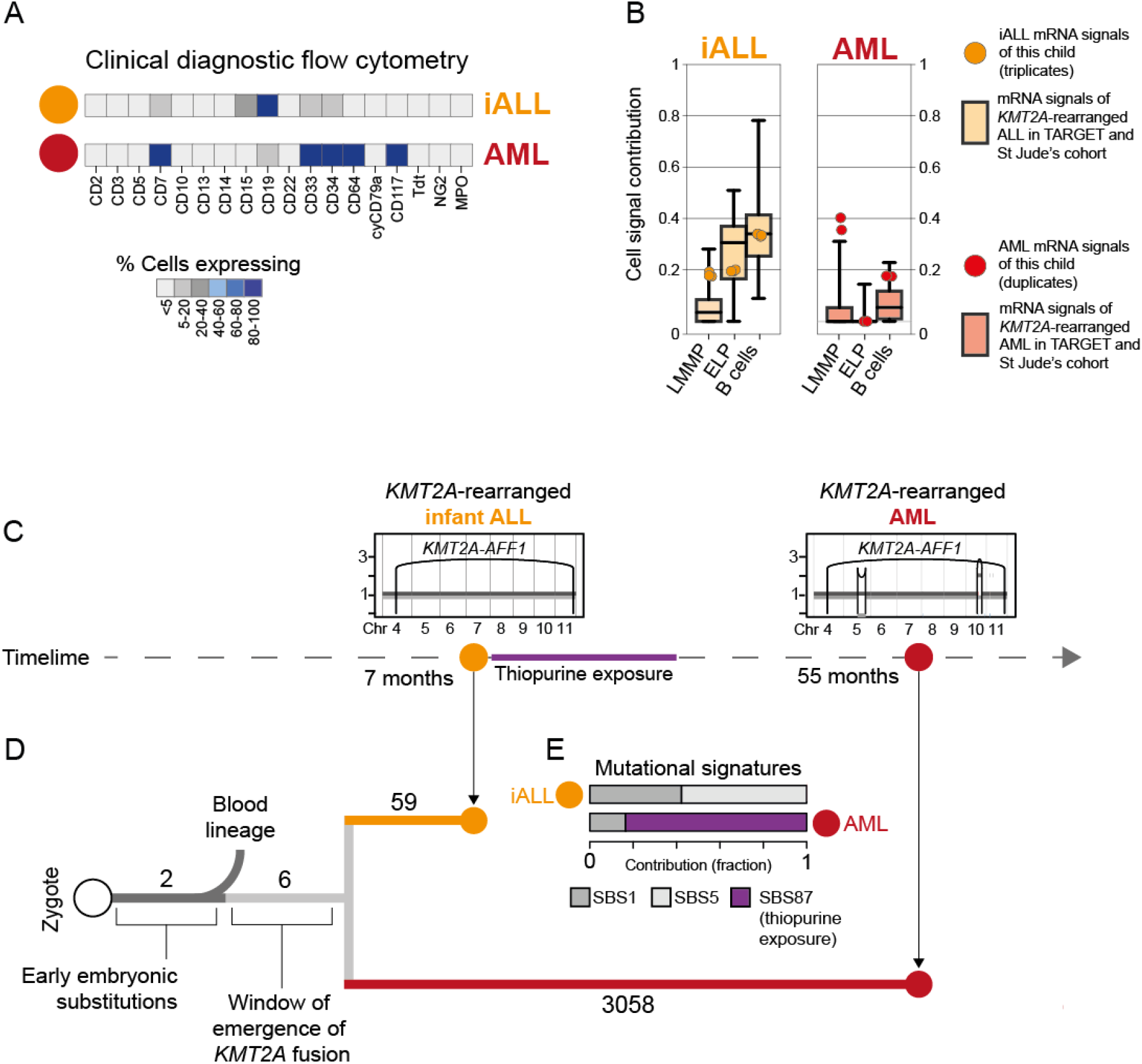
Phylogenetic timing of the origin of infant ALL. (**A**) Diagnostic flow cytometry of two leukemias that arose in the same child four years apart: *KMT2A* rearranged infant ALL (yellow, abbreviated iALL) and *KMT2A* rearranged AML (red). (B) Cell signal assessment of bulk transcriptomes generated from this child (ALL in triplicates, AML in duplicates) shows that the cell signals (LMMP; ELP; B cells, i.e. sum of all B cell signals) of ALL (yellow circle) and AML (red circle) follow the pattern of *KMT2A*-rearranged ALL (left, boxplots in background) and *KMT2A*-rearranged AML (right, boxplots in background), as defined in the St Jude’s and TARGET cohorts. Boxplot centre line=median, whiskers=min/max and box limits=25%/75% quartiles. (C) Timeline with copy number profiles of chromosomes 4 to 11 (all other chromosomes were diploid) in both leukemias showing chromosomes (x-axis) and copy number (y-axis), alleles (light and dark grey lines) as well as rearrangement breakpoints (black vertical lines and arcs) including the chromosome 4:11 translocation underpinning the *KMT2A-AFF1* fusion. (D) Phylogeny of blood and leukemia lineages with substitution burden defining each branch (number). (E) Assessment of mutational signatures as defined by the trinucleotide context of substitutions (nomenclature as per Alexandrov et al(12) highlighting in purple the dominant contribution (as % of all clonal substitutional) of signature 87 to AML. This signature is thought to be due to thiopurine agents which the child had received for ALL treatment (see timeline).

### Therapeutic hypotheses based on the ELP state of infant ALL

To distil the oncogenic features of the *KMT2A*-rearranged infant B-ALL transcriptome, we directly compared leukemia with ELP transcriptomes. We determined in independent analyses the differential gene expression between bulk *KMT2A*-rearranged infant B-ALL and published bulk ELP transcriptomes ^18^ and between single lymphoblast and single ELP cell transcriptomes (**Fig. 4A**). The overlap of these two independent analyses (**Table S6**, N=455) provided a cross-validated gene set, hereinafter referred to as the cancer core transcriptome, that differentiates *KMT2A-*rearranged B-lymphoblasts from their closest normal cell correlate (i.e. ELPs), which we annotated in four ways. Firstly, we queried whether the cancer core transcriptome contained known target genes of the *KMT2A-AFF1* fusion ^25^, the most common *KMT2A* rearrangement in B-ALL. We found 63/455 genes to be targets of the *KMT2A-AFF1* fusion, which represents a significant enrichment (p < 10^−107^, as assessed in a Monte Carlo simulation, see **Methods, Table S6**). Secondly, we asked whether the cancer core transcriptome encompassed lineage specific genes, by interrogating normal fetal bone marrow cells. We found that a subset of genes (n=51) was lineage-specific, representing either lymphoid or myeloid cell types (**Fig. 4B**). Thirdly, we annotated the cancer core transcriptome by gene ontology analysis. The top two disease annotations were lymphoblastic and myeloid leukemia, further suggesting that the cancer core transcriptome encoded a hybrid myeloid-lymphoid phenotype (**Table S6**). Fourthly, we identified cell surface antigens amongst differentially expressed genes, as many novel treatments in childhood leukemias centre around targeting blast markers through antibodies or genetically modified T-cells. 41/455 genes encoded surface markers, some of which were relatively specific to myeloid (n=18) or lymphoid (n=4) lineages, generating 72 potential non-physiological marker combinations (**Table S7**). Examples of non-physiological co-expression patterns that were particularly specific to infant B-ALL are shown in **Fig. 4C**. Interestingly, these were centred around the lymphoid marker, CD72, that a proteomic screen recently implicated as a target in infant ALL ^26^. As surface marker therapies evolve and enable the targeting of two antigens simultaneously, non-physiological co-expression of markers may represent an attractive therapeutic avenue.

**Figure 4.**
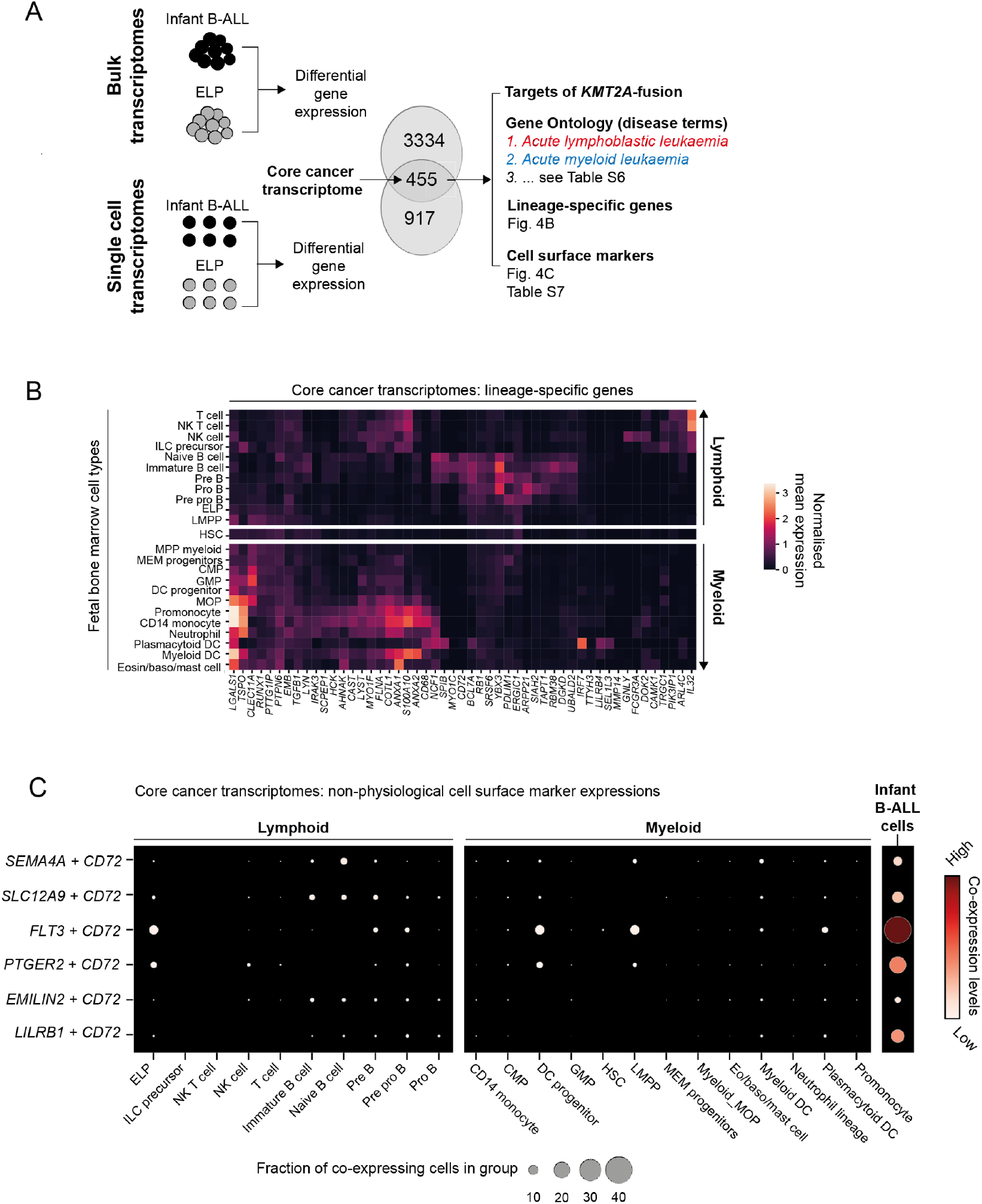
Therapeutic hypotheses based on the ELP state of infant ALL. (**A**) Distilling the core cancer transcriptome, i.e. differential gene expression between infant ALL and ELP cells from bulk and single cell data, to generate a cross-validated gene list that we annotated in three ways, as shown on right. (B) The core cancer transcriptome encodes a mixed myeloid-lymphoid phenotype. Shown is the log normalised expression (heat color) of genes (x-axis) that have relative lineage specificity in normal fetal bone marrow cell types (y-axis). (C) Examples of non-physiological combinations of cell surface markers that the core cancer transcriptome encompasses. X-axis: fetal bone marrow cell type or infant B-ALL lymphoblasts. Y-axis: marker combinations. Dots represent co-expression of the markers (average of the product of gene expression). Dot size: % of cells in cluster that express both markers; heat color: normalised co-expression level.

## Discussion

In clinical diagnostic and therapeutic terms, *KMT2A*-rearranged infant B-ALL is considered to be a B-precursor leukaemia. Based on independent data and analytical techniques, we arrived at the conclusion that infant lymphoblasts most closely resemble human fetal ELPs. This ELP-like transcriptional phenotype distinguishes *KMT2A*-rearranged infant B-ALL from other childhood B-ALL.

A key question that our findings raise is whether the ELP-like state accounts for the poor prognosis of *KMT2A*-rearranged infant B-ALL. Whilst our analyses cannot directly answer this question, three observations lend credence to this proposition. First, in both bulk mRNA and single cell analyses, *NUTM1*-rearranged infant B-ALL, recently identified to carry a favourable prognosis ^6^, exhibited cell signals away from the ELP state and more reminiscent of standard risk B-ALL. Second, we observed an ELP-like state in older children with B-ALL *KMT2A* rearrangements, in whom *KMT2A* fusions are considered a high risk cytogenetic change that mandates treatment intensification ^27^. Third, the B-ALL with the second highest relative ELP signal, B-ALL with *PAX5* alterations, is also considered high risk ^28^. These observations raise the possibility that ELP-features confer a high risk clinical phenotype in B-ALL whilst recognising the challenge of separating this signal from the prognostic significance of cytogenetic changes.

Significant efforts to identify the cell of origin in leukemias have arisen from the promise that targeted clearance of these leukaemia-initiating cells will result in durable remission. Focusing in on the cell of origin in *KMT2A*-rearranged infant B-ALL, key pieces of evidence are i) rearrangement is prenatal event, as demonstrated by Guthrie card examinations and concordance in monozygotic twins^14^, ii) rearrangement in the haematopoietic compartment is observed in CD34+CD19-cells ^29^, before VDJ recombination in most cases, resulting in low frequency of clonal immunoglobulin rearrangements ^30,31^ and iii) rearrangement may also be seen in bone marrow mesenchymal cells, suggesting a pre-haematopoietic origin in some ^31^. We directly determined the phylogenetic origin of an infant leukaemia in a rare case of a child who developed infant B-ALL and childhood AML, both harbouring *KMT2A*-rearrangements. The number of shared mutations between these leukaemias suggests that the *KMT2A*-rearrangement arose prior to gastrulation and specification of hematopoiesis. With the important caveat that this case will not represent all *KMT2A-*rearranged B-ALL cases, it demonstrates that the cell of origin does not necessarily equate with the leukemia initiating cell.

In devising novel therapeutic strategies for *KMT2A*-rearranged infant ALL, a deeper understanding of the “molecular present” of leukemic blasts may reveal more avenues for targeted therapy than deciphering their past. We compared leukemic blasts with fetal bone marrow ELPs from independent data sets to yield a core cancer transcriptome, which was characterized by fusion gene targets and a mixture of lymphoid and myeloid lineage genes. We identified non-physiological combinations of surface antigens, potentially exploitable in tandem-chimeric antigen receptor T cell or bispecific antibody therapy. Targeting combinations of antigens from different lineages simultaneously may afford exquisite specificity for cancer cells.

The quantitative molecular approach we deployed here, leveraging large archives of bulk mRNAs, emerging reference catalogues of normal human cells, and direct examination of single blast transcriptomes, lends itself for reappraising the phenotype of human leukaemias to derive novel biological and therapeutic hypotheses As leukemias are primarily classified by their hematopoietic phenotype, we propose that *KMT2A*-rearranged infant B-ALL be considered an ELP-like leukemia.

## Materials and Methods

### Ethics statement

Patient blood/bone marrow samples were obtained from the Newcastle Biobank (as approved by Newcastle & North Tyneside 1 Research Ethics Committee, reference 17/NE/0361) or GOSH diagnostic archives (as approved by National Research Ethics Service Committee London Brent, reference 16/LO/0960). Informed consent was obtained from all participants.

### Sample preparation

Peripheral blood mononuclear cells (PBMC) were prepared from blood or bone marrow by density centrifuguation, using Lymphoprep™ (Stemcell) according to manufacturer’s instructions. Samples were cryopreserved in FBS with 10% DMSO and stored in liquid nitrogen.

### Bulk RNA sequencing (lineage-switch case, Figure 3)

Total RNA was extracted from PBMC using RNeasy Mini Kit (Qiagen, Cat#74106) and mRNA captured using NEBNext Ultra Directional RNA Kit with NEBNext poly(A) mRNA Magnetic Isolation Module. Paired-end 150 bp sequencing was performed on HiSeq4000 (Illumina), with transcript abundance quantified from raw reads via Salmon v.0.8.2 and alignment performed against the hg38 reference human transcriptome (GENCODE release 27).

### Single cell RNA sequencing

Thawed cells were manually counted, and 7000 cells added to each channel of a Single Cell Chip before loading onto the 10x Chromium Controller (10x Genomics). Reverse transcription, cDNA amplification and sequencing libraries were generated using the Single Cell 3’v2 (P1_iALL, P2_iALL, P9_iAML), Single Cell 3’v3 (P3_iALL, P4_iALL, P10_iAML) and Single Cell 5’ v1 (P5_iALL, P6_iALL, P7_iALL_NUTM1 and P8_iALL_ETV6) Reagent kit (10x Genomics) according to manufacturer’s instructions. Libraries were sequenced using an Illumina HiSeq 4000 with v.4 SBS chemistry. All libraries were sequenced to achieve a minimum of 50,000 reads per cell.

### Alignment, quantification and quality control

Raw fastq files for scRNA-seq data for P1_iALL,P2_iALL, P9_iAML (single cell 3’ v2 kit) were processed with Cell Ranger v2.0.2 ^32^ pipeline and the rest of the samples were processed at later time point with Cell Ranger v3.0.2 pipeline, which aligned the reads to the reference human genome (GRCh38 v1.2.0) and produced a matrix of gene expression per single cell. Ambient mRNA contamination was removed with SoupX package v 1.4.8 in R with default parameters. Demultiplexing of P1_iALL/P2_iALL, P3_iALL/P10_iAML and P5_iALL/InfALL_classSwitch was performed with souporcell package v2.0 ^33^ with default parameters and setting number of clusters -k to 2 and --min_ref to 4 --min_alt to 4. Souporcell inferred the cluster assignment (either 0 or 1) for each single cell and given gender information (**Table S3**) we were able to demultiplex the data by checking the sex-specific gene expression in each souporcell cluster (XIST for female and RPS4Y1, ZFY and couple of others for male). Resulting gene expression matrices were further processed in python with scanpy package v1.4.4.post1 ^34^ in python and single cells were filtered to retain cells expressing >200 genes and have mitochondrial content < 20%. The code used for demultiplexing and filtering is included as a Jupyter Notebook under “Code Availability” section.

### Dimensional reduction, clustering and annotation

After filtering for low quality genes, single cell data were processed in a scanpy package in python and the total number of counts per cell were normalized to 10.000 in order to correct for library size differences; normalized data was further log-transformed. Principal component analysis (PCA) was performed on log-transformed data using default parameters (N=50), followed by computation of neighborhood graph with default parameters (N neighbors = 15) and embedding the graph in two dimensions using uniform manifold approximation and projection (UMAP). Clustering of single cell data has been performed by Louvain community detection on neighborhood graph with default resolution set to 1. Clusters were assigned as ‘cancer’ or ‘non-cancer’, based on expression of B-ALL or AML immunophenotype genes (derived from expression profiles in clinical diagnostic panels and lineage defining genes of monocytes, B cells, T cells, NK cells or progenitors, as shown in **Fig S2 and Table S4**).

### Logistic regression analysis

To measure probabilistic scores that the transcriptome of cancer cells is similar to the transcriptome of normal reference (single cell fetal bone marrow dataset) we used logistic regression as described previously ^12,16^. Briefly, a logistic regression model was trained in R using cv.glmnet function on fetal bone marrow dataset combined with Schwann cell precursor single cells from the fetal adrenal reference map ^16^, setting elastic mixing parameter alpha to 0.99 thus ensuring strong regularisation. This model was then used to predict the probabilistic score of similarity of single cells in infant leukemia dataset to cell type in fetal bone marrow dataset.

### Published bulk RNA-seq data

Pediatric tumor bulk RNA-seq data for childhood leukemia was obtained from St Jude cloud and from TARGET, together with associated metadata. Bulk RNA-seq data of human fetal bone marrow ELPs ^18^ was extracted from GEO with accession number GSE122982. Data were quantified and mapped with Salmon ^35^ with default parameters and transcript-level estimates were summarised with tximport package v 1.14.2 in R.

### Deconvolution of bulk RNAseq data

The fetal BM scRNAseq dataset was used as a reference to infer the cell type composition in bulk RNA-seq data using a previously published method of deconvolution, called cellular signal analysis ^13^. Briefly, this method aims to predict the contribution of the normal mRNA signal to each of the bulk transcriptomes. The advantage of using cellular signal analysis over other deconvolution methods is the reporting of the “unexplained signal” when the bulk transcriptome differs from all the signals in the normal reference dataset, and represented as an “Intercept” term. The model fit is based on tensorflow framework v1.14.0 and was run specifying gene weights using the geneWeights.tsv file that was supplied with the package and using default parameters for other arguments.

### Differential gene expression analysis

Differential gene expression analysis was performed using DESeq2 package v1.26.0 ^36^ in R. For bulk RNA-seq data (childhood leukemia data and ELP data) a DESeq dataset was constructed from tximport object (from Salmon quant.sf files for both childhood leukemia and ELP and creating metadata table with “group” column variables set to either ‘cancer’ or ‘ELP’). For the single cell leukemia dataset, pseudobulk was created from single cells by summarising counts per each patient. For the single cell ELP dataset, a matrix of counts was imported in Seurat and data were subsequently clustered using default parameters. Pseudobulk was created for each ELP cluster (5 in total) by summarising raw counts. Standard differential expression analysis was run using the DESeq function and the result was filtered to only include genes with adjusted p-value less than 0.05 and log2 fold changes greater than 1.

### Gene Ontology analysis

Gene ontology analysis was performed using WebGestalt (WEB-based Gene SeT AnaLysis Toolkit) ^37^. The gene list was defined as the overlap of differentially expressed genes between bulk *KMT2A*-rearranged infant B-ALL and bulk ELP transcriptomes, and between single lymphoblast and single ELP cell transcriptomes (N=455). Over-representation analysis was run using the human genome as a reference gene set and setting the disease phenotype database OMIM as a functional database.

### Analysis of enrichment of *KMT2A-AFF1* targets

Gene targets for the *KMT2A-AFF1* fusion (N=1052) were extracted from ^25^. Enrichment of these 1052 gene targets within the core leukemia transcriptome (N=455) was assessed using a Monte Carlo approach, by randomly drawing 455 genes from the possible transcriptome of 33660 genes. This step of randomly drawing the list of genes was repeated 1000 times and the p value was estimated by Student’s t-test.

### DNA sequencing and variant calling (lineage-switch case)

#### DNA sequencing and alignment

Short insert (500bp) genomic libraries were constructed and 150 base pair paired-end sequencing clusters were generated on the Illumina HiSeq XTen platform using no-PCR library protocols. DNA sequences were aligned to the GRCh37d5 reference genome by the Burrows-Wheeler algorithm (BWA-MEM) ^38^.

#### Variant calling

All classes variants were called using the extensively validated pipeline of the Wellcome Sanger Institute, built on the following algorithms: CaVEMan ^39^ for base substitutions; PINDEL for insertions/deletions ^40^; ASCAT ^41^ and *Battenberg* ^42^ for copy number changes; BRASS for structural variants ^43^.

#### Phylogenetic analyses from substitutions

We applied a previously developed framework ^44–46^. In brief, beyond the standard post-processing flags utilised in CaVEMan, we filtered out substitutions affected by mapping artefacts by setting the median alignment score of reads supporting a mutation as greater than or equal to 140 (ASMD>=140) and requiring that fewer than half of the reads were clipped (CLPM=0). Across all samples from PD38257, we recounted substitutions that were called in either blood or tumor from the patient, using a cut-off for read mapping quality (28) and base quality (25). Germline variants were removed using one-sided exact binomial test on the number of variant reads and depth present (in diploid samples) to test whether the observed counts were consistent with a true VAF of 0.5 (or 0.95 for XY chromosomes). Resulting p-values were corrected for multiple testing using the Benjamini-Hochberg method and a cut-off was set at q < 10^−5^. Variants were also filtered out if they were called in a region of consistently low or high depth in diploid regions. Variants were kept if their corresponding site had a mean depth of between 20 and 60 for autosomes and a mean depth of between 10 and 30 for the X and Y chromosome. Using a beta-binomial model of site-specific error rates as previously employed ^44–46^, we distinguished true presence of somatic variants from support due to noise. All shared substitutions were further visually inspected in the genome browser, Jbrowse ^47^. The final list substitutions included in our analyses can be found in **Table S5**.

#### Classification of SNVs

To distinguish subclonal from clonal mutations in the tumor samples, we employed a binomial mixture model to deconvolve the mutation counts into separate components. For each component, the optimal binomial probability and mixing proportion is estimated using an expectation-maximisation algorithm. The optimal number of components is determined by the Bayesian information criterion. If the binomial probability of a component approximates the expected VAF (0.5 for diploid regions) adjusted for tumor purity, the mutations assigned to that cluster are classified as clonal. If the estimated binomial probability for a component is lower, it is classified as subclonal.

#### Mutational signature analysis

Mutation signatures were fitted to the trinucleotide counts of SNVs in the main clone and subclone of the ALL (PD38527a) and the AML (PD38527c) using the SigFit algorithm ^48^ and the COSMIC reference database of mutational signatures (https://cancer.sanger.ac.uk/signatures/sbs/, V3.2), as employed previously ^49^. Initially, all reference signatures were fitted to the mutation counts. Only signatures that contributed at least 2% were retained during the subsequent fitting. Where mutation counts were low (<100), erroneous C>T signatures, such as those from UV exposure (SBS7a) or mismatch repair deficiency (SBS6), were attributed to the samples. Because of their biological implausibility, these signatures were removed from the final set of fitted signatures.

#### Data and code availability

Jupyter notebook for processing single cell data, including cellranger filtered counts data and steps to reproduce Figure 1 and 2, is available at https://github.com/kheleon/leukemia-paper.

## Acknowledgments

We thank Irene Roberts and Anindita Roy, both from the University of Oxford, and Aidan Maartens (science writer at the Wellcome Sanger Institute), for critical review of the manuscript. We are indebted to patients and their families for participating in our research.

## Funding

Wellcome Trust WT206194 (Wellcome Sanger Institute); WT110104/Z/15/Z (SBehjati); WT107931/Z/15/Z (MH)

Lister Institute of Preventative Medicine (MH)

Newcastle NIHR-Biomedical Research Centre (MH and LJ)

National Institute for Health Research Academic Clinical Lectureship (LJ)

Medical Research Council MRC clinician scientist fellowship MR/S021590/1 (SBomken) CRUK program grant (C27943/A12788) (OH)

Kay Kendall Leukaemia Fund (KKL1142) (OH) Kika program grant (329) (OH)

Children with Cancer UK 14-169,17-249; NIHR Great Ormond Street Hospital Biomedical Research Centre (OW)

## Author contributions

Conceptualization: SBehjati, JB

Methodology: SBehjati, EK, MDY, SW, THHC

Investigation: EK, LJ, TDT, THHC, JE, TP, EP, KS

Visualization: SBehjati, EK, LJ

Funding acquisition: SBehjati

Supervision: SBehjati, JB, MH, SBomken

Writing – original draft: SBehjati, EK, LJ

Writing – review & editing: SBomken, MH, GC, ST, OW, OH

## Data and materials availability

Raw sequencing data that we generated have been deposited as follows: DNA sequences of the lineage switch case (PD38257a to c) have been deposited in the European Genome-phenome Archive under study ID EGAD00001007853; corresponding RNA sequences have been deposited in the NCBI Sequence Read Archive under project IDs PRJNA547947 and PRJNA547815; single cell RNA sequences have been deposited in the European Nucleotide Archive (accession number ERP125305) and in the European Genome-phenome Archive (accession number EGAD00001007854).

## Supplementary Materials

Figs. S1-S2

Table S1. Categories of childhood leukemia from St Jude and TARGET cohorts

Table S2. Mapping of fetal bone marrow reference populations to detailed cell_id in fetal bone marrow dataset

Table S3. Case details for infant leukemia samples

Table S4. Immunophenotypes of infant leukemia samples

Table S5. List of single nucleotide variants in lineage-switch case analysis

Table S6. Gene ontology annotation of differentially expressed genes between infant ALL and ELP cells from bulk and single-cell mRNA data

Table S7. Surface markers from list of differentially expressed genes between infant ALL and ELP cells from bulk and single-cell mRNA data

