## Supplementary Information for "Single cell mRNA signals reveal a distinct developmental state of *KMT2A*-rearranged infant B-cell acute lymphoblastic leukemia"

**This PDF file includes:**

Figs. S1 to S2
Tables S1 to S7 (S5 and S6 as separate files)

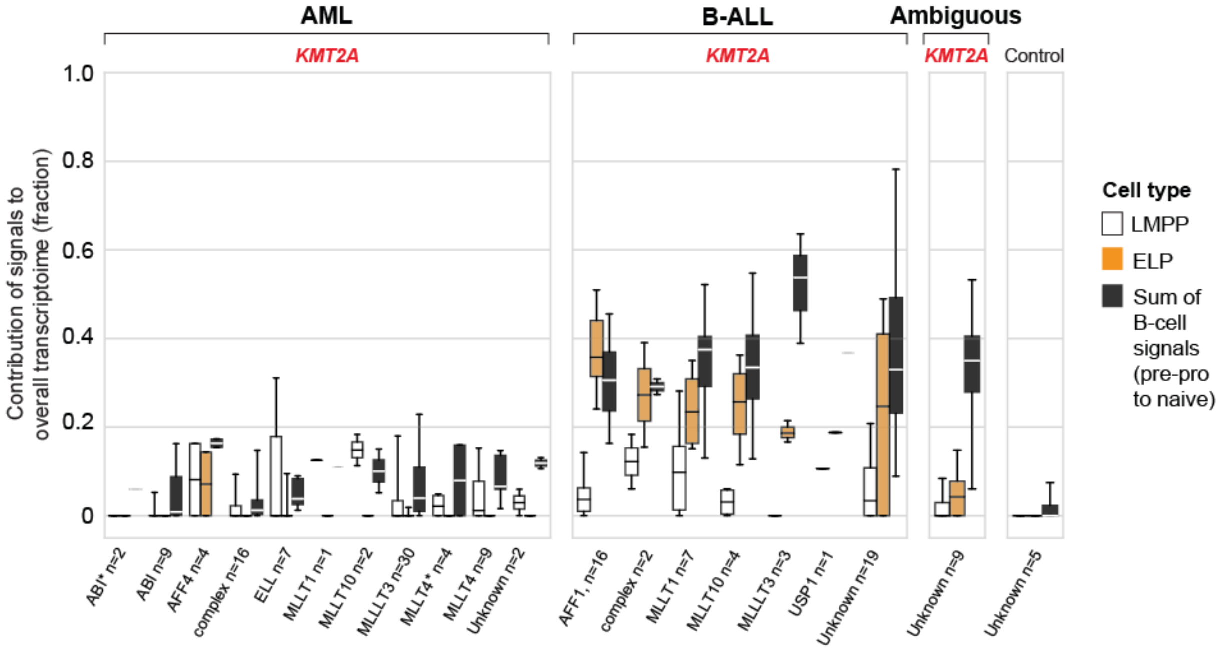

**Fig. S1. ELP signal is common to *KMT2A* rearranged B-ALL, independent of fusion partner**

Box and whisker plots showing contributions of cell signals – LMPP, ELP and latter B-cell stages (i.e. pre-pro B, pro-B, pre-B and naive B combined) to the transcriptome of *KMT2A*-rearranged leukemias grouped by *KMT2A* fusion partner (see x-axis labels).

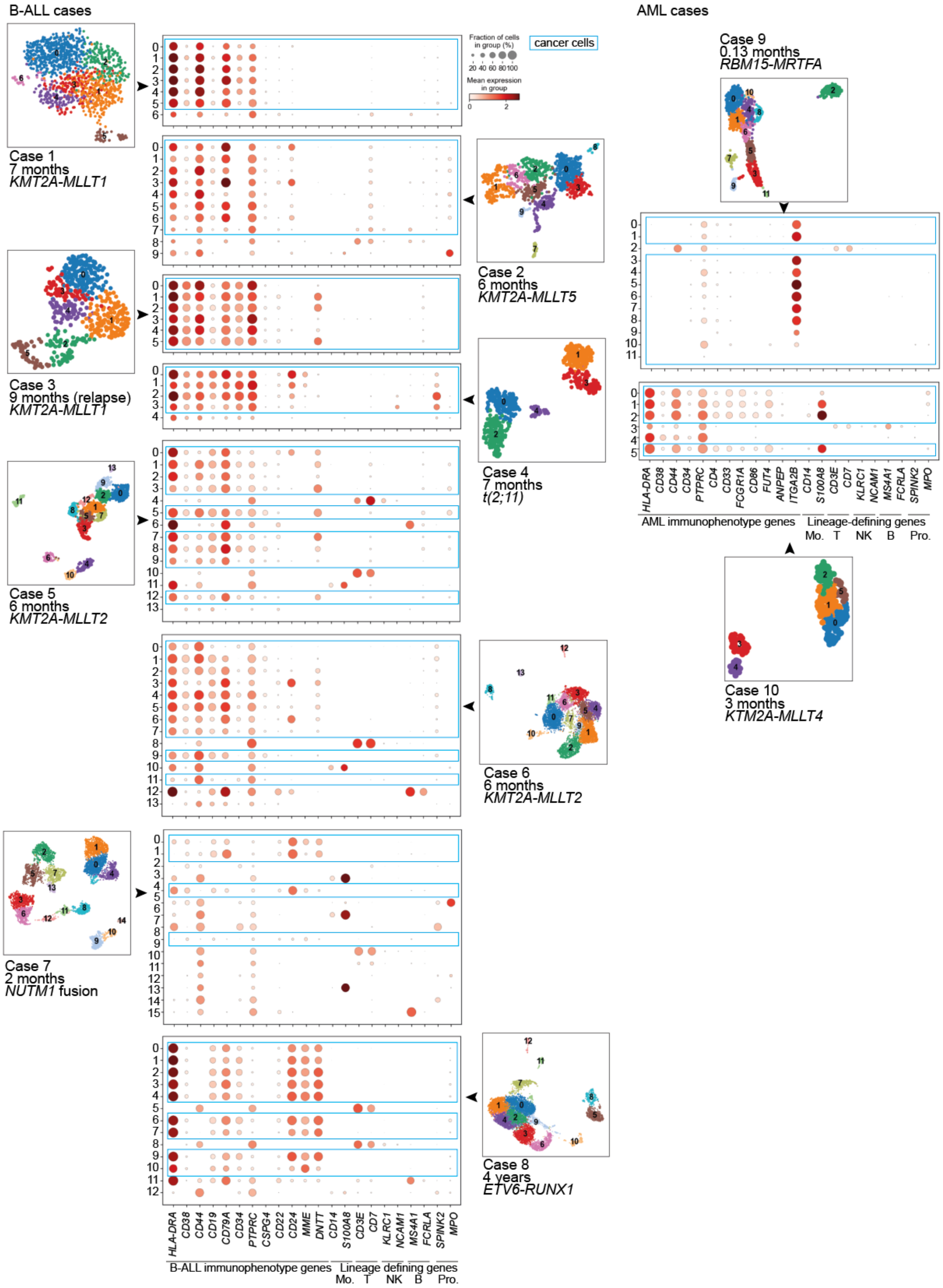

B-ALL cases
AML cases
Fraction of cells in group (%)
20 40 60 80 100
Mean expression in group
0 2
cancer cells
Case 1
7 months
KMT2A-MLLT1
Case 2
6 months
KMT2A-MLLT5
Case 3
9 months (relapse)
KMT2A-MLLT1
Case 4
7 months
t(2;11)
Case 5
6 months
KMT2A-MLLT2
Case 6
6 months
KMT2A-MLLT2
Case 7
2 months
NUTM1 fusion
Case 8
4 years
ETV6-RUNX1
Case 9
0.13 months
RBM15-MRTFA
Case 10
3 months
KTM2A-MLLT4
HLA-DRA CD38 CD44 CD19 CD79A CD34 PTPRC CSPG4 CD22 CD24 MME DNTT CD14 S100A8 CD3E CD7 KLRC1 NCAM1 MS4A1 FCRLA SPINK2 MPO
B-ALL immunophenotype genes
Lineage defining genes
Mo. T NK B Pro.
HLA-DRA CD38 CD44 CD34 PTPRC CD4 CD33 FCGR1A CD86 FUT4 ANPEP ITGA2B CD14 S100A8 CD3E CD7 KLRC1 NCAM1 MS4A1 FCRLA SPINK2 MPO
AML immunophenotype genes
Lineage-defining genes
Mo. T NK B Pro.

**Fig. S2. Identification of cancer cells in leukemia scRNAseq data using immunophenotype gene expression**

UMAP projections of leukaemia scRNA-seq data sets, coloured by Louvain cluster. Accompanying dotplots show per-cluster expression of B-ALL immunophenotype genes or AML immunophenotype genes and lineage-defining genes of monocytes (Mo.), T cells (T), NK cells (NK), B cells (B) and progenitors (Pro.). Dot colour denotes log-transformed, normalised and scaled gene expression value, while dot size indicates percentage of cells in each cluster expressing the stated gene. Immunophenotypes are provided in **Table S4**.

| Diagnosis | Cohort | Age group | Molecular/cytogenetic abnormality | Number of cases | Categories merged |
|---|---|---|---|---|---|
| B-ALL | St Jude | child | iAMP21 | n=16 | |
| B-ALL | St Jude | child | ZNF384 rearrangement | n=5 | ZNF384 rearrangement (n=3), ZNF384 rearrangement-like (n=2) |
| B-ALL | St Jude | child | TCF3-PBX1 | n=10 | |
| B-ALL | TARGET | child | TCF3-PBX1 | n=11 | |
| B-ALL | St Jude | child | PAX5 | n=22 | PAX5 alteration (n=16), PAX5 P80R (n=6) |
| B-ALL | TARGET | child | NOS | n=71 | |
| B-ALL | St Jude | child | NOS | n=17 | |
| B-ALL | St Jude | child | MEF2D rearrangement | n=6 | |
| B-ALL | St Jude | child | IGH-CEBPD | n=2 | |
| B-ALL | St Jude | child | Hypodiploidy | n=8 | |
| B-ALL | St Jude | child | Hyperdiploidy | n=37 | |
| B-ALL | TARGET | child | Hyperdiploidy | n=20 | |
| B-ALL | St Jude | child | ETV6-RUNX1 | n=72 | ETV6-RUNX1 (n=68), ETV6-RUNX1-like (n=4) |
| B-ALL | TARGET | child | ETV6-RUNX1 | n=5 | |
| B-ALL | St Jude | child | DUX4-IGH | n=58 | DUX4-IGH (n=57) and DUX4-IGH-like (n=1) |
| B-ALL | TARGET | child | BCR-ABL1 | n=5 | |
| B-ALL | St Jude | child | BCR-ABL1 | n=34 | |
| B-ALL | St Jude | child | BCR-ABL1 like | n=52 | |
| B-ALL | TARGET | child | KMT2A rearrangement | n=4 | |
| B-ALL | St Jude | child | KMT2A rearrangement | n=12 | |
| B-ALL | St Jude | infant | NUTM1 rearrangement | n=2 | |
| B-ALL | St Jude | infant | KMT2A rearrangement | n=36 | |
| ALAL | St Jude | child | NOS | n=3 | |
| ALAL | TARGET | child | NOS | n=106 | |
| ALAL | TARGET | child | BCR-ABL1 | n=2 | |
| ALAL | TARGET | child | KMT2A rearrangement | n=2 | |
| ALAL | TARGET | infant | NOS | n=4 | |
| ALAL | TARGET | infant | KMT2A rearrangement | n=7 | |
| AML | St Jude | child | Promyelocytic | n=3 | |
| AML | TARGET | child | NOS | n=476 | |
| AML | St Jude | child | NOS | n=26 | |
| AML | St Jude | child | Core Binding Factor | n=35 | |
| AML | St Jude | child | KMT2A rearrangement | n=9 | |
| AML | TARGET | child | KMT2A rearrangement | n=60 | |
| AML | TARGET | infant | NOS | n=21 | |
| AML | St Jude | infant | NOS | n=2 | |
| AML | TARGET | infant | KMT2A rearrangement | n=17 | |
| AMKL | St Jude | child | NOS | n=96 | |
| AMKL | St Jude | infant | NOS | n=7 | |
| T-ALL | TARGET | child | NOS | n=265 | |
| T-ALL | St Jude | child | NOS | n=15 | |
| T-ALL | St Jude | child | KMT2A rearrangement | n=2 | |
| T-ALL | St Jude | infant | NOS | n=2 | |

**Table S1. Categories of childhood leukemia from St Jude and TARGET cohorts**

| | | | | | | | | | |
|---|---|---|---|---|---|---|---|---|---|
| | Shown in Fig. 1B | | | | | | | | |
| BM reference states | LMPP | ELP | Pre pro B | Pro B | Pre B | Immature B | Naïve B | NK | T |
| Cell_id in Jardine et al. | LMPP | ELP | Pre pro B | Pro B | Pre B | Immature B | Naïve B | CD56 bright NK | CD4 T cell |
| | | | | | | | | mature NK | CD8 T cell |
| | | | | | | | | | Treg |
| | Shown in Fig. 1B ctd | | | | | | | | |
| BM reference states | MPP myeloid | CMP | MOP | Promono. | CD14 mono. | Neut. Lineage | MEM progenitors | MK | Eo/baso/mast |
| Cell_id in Jardine et al. | MPP myeloid | CMP | MOP | Promonocyte | CD14 monocyte | promyelocyte | eo/baso/mast precursor | early MK | eosinophil |
| | | | | | | myelocyte | MEMP | MK | basophil |
| | | | | | | metamyelocyte | MEP | | mast cell |
| | | | | | | neutrophil | | | |
| | Not shown in Fig. 1B (low deconvolution signal) | | | | | | | | |
| BM reference states | non-haematopoietic | pDC | myeloid DC | macrophage | HSC | GMP | erythroid | DC progenitor | |
| Cell_id in Jardine et al. | arteriolar fibroblast | plasmacytoid DC | DC1 | stromal macrophage | HSC | GMP | early erythroid | myeloid DC progenitor | |
| | chondrocyte | | DC2 | monocytoid macrophage | | | mid erythroid | pDC progenitor | |
| | early osteoblast | | DC3 | erythroid macrophage | | | late erythroid | | |
| | endosteal fibroblast | | tDC | | | | | | |
| | immature EC | | | | | | | | |
| | muscle | | | | | | | | |
| | muscle stem cell | | | | | | | | |
| | myofibroblast | | | | | | | | |
| | osteoblast | | | | | | | | |
| | osteoblast precursor | | | | | | | | |
| | osteochondral precursor | | | | | | | | |
| | osteoclast | | | | | | | | |
| | proliferating EC | | | | | | | | |
| | schwann cells | | | | | | | | |
| | sinusoidal EC | | | | | | | | |
| | tip EC | | | | | | | | |

**Table S2. Mapping of fetal bone marrow reference populations to detailed cell_id in fetal bone marrow dataset (*9*)**

| Case ID | Age (months) | Sex | Disease | Cytogenetics | Timepoint | Cells | Experiment | PR ID | PD ID | Sc ID |
|---|---|---|---|---|---|---|---|---|---|---|
| | | | | | Diagnosis | NA | BulkRNA | PR40839a | | |
| | 7 | Male | Infant ALL | (q21;q23) aka KMT2A-MLLT2 | Diagnosis | NA | BulkRNA | PR40839c | | |
| | | | | | Diagnosis | NA | BulkRNA | PR40839d | | |
| Lineage-switch | | | | | Diagnosis | NA | BulkDNA | | PD38257a | |
| | | | | | Relapse | NA | BulkRNA | PR40839f | | |
| | 55 | Male | Infant AML | t(4;11)(q21;23)/ 46, idem, del(5)(q2?q3?, add(10)(q)/ 47, idem, +der(4) t(4;11) | Relapse | NA | BulkRNA | PR40839g | | |
| Lineage-switch | | | | | Relapse | NA | BulkDNA | | PD38257c | |
| Lineage-switch | 56 | Male | Remission | not applicable | Remission | NA | BulkDNA | | PD38257b | |
| 1 | 7 | Male | Infant ALL | t(11;19)(q23;p13.3) and t(13;17)(q1?2;q21) aka KMT2A-MLLT1 | Diagnosis | 1274 | 10X | PR43971a | PD43971a | 4602STDY7920960+4602STDY7920961 |
| 2 | 6 | Female | Infant ALL | t(1;11)(p32;q23) aka KMT2A-MLLT5 | Diagnosis | 1047 | 10X | PR43972a | PD43972a | 4602STDY7920960+4602STDY7920961 |
| 3 | 9 | Female | Infant ALL | t(11;19) aka KMT2A-MLLT1 | Relapse | 541 | 10X | | | CG_SB_NB8791864 |
| 4 | 7 | Female | Infant ALL | t(2;11) and Trisomy 8 | Diagnosis | 616 | 10X | | | CG_SB_NB8791866+CG_SB_NB8791867 |
| 5 | 6 | Female | Infant ALL | t(4;11)(q21;q23) aka KMT2A-MLLT2 | Diagnosis | 1878 | 10X | | | WSSS_F_BON10188628 |
| 6 | 5 | Female | Infant ALL | t(4;11)(q21;q23) aka KMT2A-MLLT2 | Diagnosis | 7141 | 10X | | | WSSS_F_BON10188629 |
| 7 | 2 | Male | Infant ALL | ACIN1-NUMT1 fusion | Diagnosis | 6017 | 10X | | | WSSS_F_BON10188631 |
| 8 | 48 | Male | Infant ALL | ETV6-RUNX1 fusion | Diagnosis | 8474 | 10X | | | WSSS_F_BON10188630 |
| 9 | 0 | Female | Infant AML | t(1;22)(p13;q13) | Diagnosis | 1149 | 10X | PR43973a | PD43973a | 4602STDY7920965 |
| 10 | 3 | Male | Infant AML | t(6;11) aka KMT2A-MLLT4 | Diagnosis | 297 | 10X | | | CG_SB_NB8791864 |

**Table S3. Case details for infant leukemia samples.**
Case IDs indicate how samples are referenced in manuscript, figures and supplementary materials.  PR, PD, and Sc IDs are Sanger Institute references for bulk RNA-seq, DNA-seq and single-cell RNA-seq data.

| | Antigen | Case 1 | Case 2 | Case 3 | Case 4 | Case 5 | Case 6 | Case 7 | Case 8 | Case 9 | Case 10 | Gene name |
|---|---|---|---|---|---|---|---|---|---|---|---|---|
| Surface | CD10 | 0 | 0 | 0 | 15 | - | - | 96 | + | 0 | 0 | *MME* |
| | CD117 | 0 | 0 | 0 | 0 | NT | NT | 0 | NT | 21 | 0 | *KIT* |
| | CD11b | NT | NT | NT | 0 | NT | NT | NT | NT | 0 | 54 | *ITGAM* |
| | CD13 | 16 | 6 | 0 | 11 | NT | NT | 44 | weak | 61 | 44 | *ANPEP* |
| | CD14 | NT | NT | NT | 0 | NT | NT | NT | NT | 0 | 16 | *CD14* |
| | CD15 | NT | NT | NT | 49 | + | NT | NT | NT | 0 | 95 | *FUT4* |
| | CD16 | 0 | 0 | 0 | 0 | NT | NT | 0 | NT | 0 | 0 | *FCGR3A* |
| | CD19 | 99.5 | 100 | 100 | 100 | + | + | 100 | + | 0 | 0 | *CD19* |
| | CD2 | 0 | 0 | 0 | 0 | NT | NT | 0 | NT | 71 | 84 | *CD2* |
| | CD20 | 0 | 0 | 0 | 0 | - | NT | 0 | - | 0 | 0 | *MS4A1* |
| | CD22 | 81 | 27 | 95 | 15 | + | NT | 96 | + | 0 | NT | *CD22* |
| | CD24 | 12.5 | 65 | 4 | 59 | NT | NT | 100 | NT | 0 | NT | *CD24* |
| | CD3 | 0 | 0 | 0 | 0 | NT | NT | 0 | NT | 0 | 0 | *CD3E* |
| | CD33 | 24 | 0 | 0 | 5 | NT | - | 21 | + | 78 | 100 | *CD33* |
| | CD34 | 96 | 4 | 98 | 100 | +/- | +/- | 0 | + | 44 | 100 | *CD34* |
| | CD38 | 100 | 100 | 100 | 100 | + | NT | 100 | NT | 20 | 100 | *CD38* |
| | CD4 | 0 | 0 | 0 | 0 | NT | NT | 0 | NT | 100 | 100 | *CD4* |
| | CD41a | NT | NT | NT | NT | NT | NT | NT | NT | 100 | 0 | *ITGA2B* |
| | CD42b | NT | NT | NT | NT | NT | NT | NT | NT | 100 | 0 | *GPB1* |
| | CD44 | 100 | 100 | 100 | 100 | NT | NT | 10 | NT | 0 | 100 | *CD44* |
| | CD45 | 100 | 100 | 100 | 100 | NT | NT | 61 | NT | 100 | 100 | *PTPRC* |
| | CD5 | 0 | 0 | 0 | 0 | NT | NT | 0 | NT | 0 | 0 | *CD5* |
| | CD61 | NT | NT | NT | NT | NT | NT | NT | NT | 100 | 0 | *ITGB3* |
| | CD64 | NT | NT | NT | 2 | NT | NT | NT | NT | NT | 100 | *FCGR1A* |
| | CD66 | 0 | 0 | NT | NT | NT | | NT | NT | NT | NT | *CEACAM1* |
| | CD7 | 0 | 67 | 0 | NT | NT | NT | 0 | NT | 33 | 0 | *CD7* |
| | CD79b | 0 | 0 | 0 | 0 | NT | NT | 0 | NT | 0 | 0 | *CD79B* |
| | CD8 | 0 | 0 | 0 | 0 | NT | NT | 0 | NT | 0 | 0 | *CD8A* |
| | CD86 | 22 | 12 | 0 | 67 | NT | NT | 0 | NT | 0 | 90 | *CD86* |
| | HLA-DR | 100 | 94 | 100 | 100 | + | + | 100 | + | 0 | 100 | *HLA-DRA* |
| | NG2 | 81 | 78 | 95 | 14 | NT | NT | 0 | NT | 0 | 41 | *CSPG4* |
| | Surface Ig | NT | NT | NT | NT | - | NT | NT | - | NT | NT | |
| Intracellular | Cy3 | 3 | 3 | 0 | 4 | NT | NT | NT | NT | NT | NT | *CD3E* |
| | Cy79a | 95 | 88 | 100 | 83 | NT | low | 100 | NT | NT | NT | *CD79A* |
| | MPO | 0 | 6 | 0 | 3 | NT | NT | NT | NT | NT | NT | *MPO* |
| | TdT | 0 | 18 | 65 | 24 | + | - | 100 | + | NT | NT | *DNTT* |

Genes shown in Fig. S2
- B-ALL immunophenotype genes (blue)
- AML immunophenotype genes (yellow)
- B-ALL and AML immunophenotype genes (green)

**Table S4. Immunophenotypes of infant leukemia samples**
Diagnostic immunophenotypes of leukemia samples. Antigen selection and reporting was performed according to local clinical practice. NT= not tested. Numerical data indicates % of leukemia cells expressing given antigen. + indicates expressed by leukaemia cells; - indicates not expressed and +/- indicates expressed by a proportion of leukemia cells. Row colors denote immunophenotype genes shown in **Fig. S2**.

**Table S5 (separate file). List of single nucleotide variants in lineage-switch case analysis**

**Table S6 (separate file). Gene ontology annotation of differentially expressed genes between infant ALL and ELP cells from bulk and single cell mRNA data.**

| Gene differentially expressed between ELP and infant B-ALL | Myeloid (M), B-lymphoid (B) or lineage non-specific (NS)* |
|---|---|
| ADAM10 | NS |
| CD68 | M |
| CD70 | NS |
| CD72 | B |
| CD86 | M |
| CPD | M |
| CSPG4 | NS |
| EMB | M |
| EMILIN2 | M |
| FCGR2B | M |
| FCGR3A | M |
| FGFRL1 | NS |
| FLT3 | M |
| GREM1 | NS |
| HEG1 | M |
| HLA-B | NS |
| HLA-F | B |
| ICOSLG | B |
| IL10RA | M |
| ITGAM | M |
| LAMP1 | NS |
| LILRB1 | M |
| LRP5 | NS |
| LTB4R | M |
| LTBP3 | NS |
| MUC4 | NS |
| PLXNA3 | NS |
| PLXNB1 | NS |
| PTGER2 | M |
| PTPRC | NS |
| PTTG1IP | M |
| QSOX2 | NS |
| SEMA4A | M |
| SIDT2 | NS |
| SLC12A9 | M |
| SPPL2B | NS |
| TGFB1 | M |
| TGFBR2 | NS |
| TTYH3 | NS |

*Myeloid or B lymphoid annotation based on % cells expression gene and expression level in fetal BM B lineage (B)> myeloid lineage, vice versa (M)

**Table S7. Surface markers from list of differentially expressed genes between infant ALL and ELP cells from bulk and single cell mRNA data.**
